# CTCF regulates wild-type and recombinant AAV gene expression by shaping viral chromatin

**DOI:** 10.64898/2026.06.03.729793

**Authors:** Clairine I.S. Larsen, Rhiannon R. Abrahams, Rea Guertler, Elliott M. Wilion, Eda Erata, Zachary H. Sykes, Lorelei Stoica, Gopishankar Thirumoorthy, Divya Sinha, Richa Rai, Elizabeth K. Hofer, Eleanor Hart, David M. Gamm, Matthew S. Fuller, Kinjal Majumder

## Abstract

Adeno-Associated Viruses (AAVs) are powerful platforms for delivering therapeutic transgenes via recombinant AAV (rAAV) vectors. However, a limited understanding of the regulation of AAV gene expression has narrowed the ability to efficiently express therapeutic transgenes from rAAV vectors. Since rAAVs retain only the wtAAV inverted terminal repeats (ITR), we hypothesized that regulatory elements outside the ITR that govern wild-type AAV (wtAAV) gene expression can be used to modify rAAV genomes to enhance vector performance. Through in silico analysis, biochemical pulldowns, and high-throughput sequencing, we have identified that the host architectural protein CCCTC-binding Factor (CTCF) associates with the wtAAV type 2 (wtAAV2) genome but is absent from rAAV vectors. Global knockdown and site-specific deletion revealed that the CTCF binding element (CBE) on the wtAAV2 genome, located upstream of the viral P5 promoter, regulates expression of the viral *Rep68/78* genes. We have re-engineered new rAAV vectors expressing a GFP reporter transgene to contain the wtAAV2-CBE upstream of the vector promoter. Our results show that CTCF binding dramatically increased rAAV transduction efficiency and GFP expression by up to four-fold across multiple cell types. This enhancement was independent of the AAV capsid serotype used for packaging rAAV vectors. CUT&RUN analysis revealed that this CBE was necessary and sufficient to regulate the chromatin landscape of wtAAV2 and rAAV2. Finally, we observed that CTCF-mediated chromatin remodeling of rAAV2 led to increased production of nascent RNA transcripts from the vector genome. Based on our findings, we propose that CTCF supports wtAAV2/rAAV gene expression by shaping the local chromatin landscape.

**SIMPLE ABSTRACT:** Recombinant Adeno-Associated Viruses (rAAV) gene therapy vectors have been engineered from wild-type AAV (wtAAV) by inserting the viral telomeres (that serve as replication and packaging signals) on either side of therapeutic transgenes. However, efficient expression of transgenes using current rAAV technologies require high doses, which can lead to sporadic toxic side effects. We hypothesized that uncharacterized regulatory elements in the wtAAV2 genome drive efficient viral gene expression and are absent from the current generation of rAAV vectors. Using in-silico analysis combined with biochemical pulldowns, high-throughput sequencing, and mutant viral systems, we have identified a novel *cis*-acting element bound by the cellular architectural protein CCCTC-binding Factor (CTCF). This CTCF binding element is necessary for wtAAV2 gene expression and is sufficient to enhance the rAAV vector’s ability to express reporter transgenes. This CTCF-binding element regulates the chromatin landscape of the virus and its vectors. Our discovery that adding the 19-bp AAV2 CTCF-binding element enhances transgene expression without affecting vector production efficiency presents a promising new rAAV gene therapy platform that is likely to reduce clinical doses and minimize toxicity in therapeutic applications.

## INTRODUCTION

Wild type Adeno-Associated Virus Type 2 (wtAAV2) is a small single-stranded DNA (ssDNA) virus of the family *Parvoviridae* that is utilized in gene therapy to deliver therapeutic transgenes as recombinant AAV (rAAV) vectors[1]. The wtAAV2 capsid delivers the 4.7 kb ssDNA genome to the nucleus, where it is processed to generate a double-stranded DNA molecule that serves as the platform for expressing viral (wtAAV2) or therapeutic (rAAV) transgenes[2, 3]. The wtAAV2 genome is flanked by palindromic sequences that generate hairpin-like structures known as Inverted Terminal Repeats (ITRs; [4]). These ITRs are the viral telomeres that serve as the origin of replication and the packaging signal [5, 6]. wtAAV2 has evolved multiple strategies to compress its genome, including overlapping open reading frames [encoding distinct isoforms of non-structural replication (Rep) proteins (Rep40/52/68/78) and capsid (Cap) proteins (VP1/2/3)], alternative splicing, and multiple compact transcriptional regulatory elements[7–11]. Recent studies have identified two additional accessory proteins, Assembly-Activating Protein (AAP; [12, 13]) and Membrane-Associated Accessory Protein (MAAP; [14, 15]), that regulate the viral life cycle by serving as a chaperone protein and a cellular egress factor, respectively. These wtAAV2 proteins interact with partners in the host milieu that are critical for driving the outcome of viral infection.

As a member of the *Dependoparvovirus* genus, wtAAV2 requires a coinfecting helper virus (adenovirus[16, 17], herpes-simplex virus[18, 19], vaccinia virus[20], human papillomavirus[21], or human bocavirus 1[22]) to efficiently express viral genes, replicate the viral genome, package and produce infectious progeny [23]. In the absence of a coinfecting virus (termed mono-infection), wtAAV2 expresses low levels of the transcripts of its non-structural Rep proteins. This ability to express an open reading frame located between the ITRs in the absence of viral replication is leveraged to use wtAAV2 as a gene therapy platform [1]. rAAVs are engineered to retain only the wtAAV2 ITRs, with all viral open-reading frames and non-ITR-associated regulatory elements replaced by a therapeutic transgene and exogenous regulatory elements. rAAV vectors are primarily produced by co-transfection of plasmids expressing the vector genome, the essential proteins from adenovirus (AdV) required for replication and packaging (E1, E2a, E4orf6, and VA RNA), and the wtAAV2 replication and capsid genes (Rep-Cap)[16, 17, 24–27]. This co-transfection generates encapsidated rAAV particles that can be delivered to target cells and express the therapeutic transgene[28, 29]. While rAAV-based gene therapies have shown clinical success, with seven gene therapies being approved by the FDA[1], the requirement for extremely high dosages (up to 10^14^ vector genomes per kilogram in patients[30]) suggests that a knowledge gap exists in being able to efficiently express therapeutic transgenes using rAAV technology. This necessitates a better understanding of how wtAAV2 genes are expressed.

Transcription is topologically regulated by spatial interactions of distally bound enhancers and promoters impacting local chromatin accessibility [31, 32]. Indeed, wtAAV2 was one of the first systems in which the phenomenon of transactivation of viral promoters by looping out intervening DNA sequences was observed[33–35]. Extensive characterizations of mammalian genome topology have revealed that the cellular protein CCCTC-binding factor (CTCF) plays a central role in maintaining the three-dimensional genome structure and in regulating transcription by bringing distally located promoters and enhancers into spatial proximity [31, 36, 37]. To facilitate these interactions, dimerized CTCF serves as an anchor at the base of the genome loop and is stabilized by the Cohesin complex (composed of SMCs and RAD21 subunits), extruding the intervening DNA [38–40]. CTCF also functions as a chromatin insulator, demarcating the boundary between active and inactive chromatin [41]. Importantly, the functions of CTCF have been widely usurped by viral pathogens to control viral gene expression and life cycle[42, 43]. The gammaherpesviruses Epstein-Barr Virus (EBV) and Kaposi’s Sarcoma-associated Herpesvirus (KSHV) possess CTCF-binding sites at promoters and control regions that generate 3D loops on the viral genome, regulating the stages of viral latency and the latent-to-lytic switch [44–49]. In the small DNA virus Human Papillomavirus (HPV), CTCF interacts with the transcription factor Yin-Yang 1 (YY1) to bring together distal regions of the viral genome, generating a 3D loop specific to high-risk HPVs that regulates viral oncogene expression [50, 51]. During differentiation-dependent amplification of HPV, the CTCF-interacting protein SMC1, a subunit of cellular Cohesin complex, aids in viral genome amplification[52]. Independent of architectural proteins, CTCF regulates viral and cellular gene expression by directly interacting with the large subunit of RNA Pol II[53], directing RNA Pol II pausing[54], elongation [55], and alternative splicing[54, 56]. These mechanisms of CTCF-mediated control of RNA production and processing have been observed in the autonomous parvovirus Minute Virus of Mice (MVM; [57]) and in Hepatitis B Virus (HBV; [58]). The widespread usage of CTCF by viruses highlights its importance in regulating gene expression.

CTCF has been utilized to enhance the efficiency of other viral-based gene therapy platforms. In the AdV platform, the insulator function of CTCF was leveraged to suppress viral genes near the exogenous transgene promoter while driving efficient transgene expression. As a result, the innate immune response to the vector was attenuated [59]. CTCF binding has been utilized to enhance the production of rAAV vectors. The addition of CTCF-binding elements in the bacterial backbone of rAAV infectious clone reduced deleterious cross-packaging of the plasmid backbone into AAV capsids, increasing the production of correctly encapsidated vector genomes[60]. Together, these applications establish CTCF as a versatile tool for enhancing viral gene therapy platforms.

In this study, we have discovered a role for CTCF in regulating wtAAV2 gene expression and engineered an rAAV vector with a CTCF-binding site to enhance transgene expression. wtAAV2 contains a native CTCF binding element (AAV2-CBE) upstream of the viral promoter P5. Global CTCF knockdown and deletion of the AAV2-CBE reduced Rep68/78 transcripts. Insertion of the AAV2-CBE upstream of the CMV promoter in an rAAV vector expressing a GFP reporter transgene led to enhanced GFP expression across multiple cell types delivered by multiple capsid serotypes. We show that the increase in viral and vector expression is associated with CTCF-mediated changes in the wtAAV2/rAAV chromatin landscape, leading to enhanced nascent RNA transcription. Taken together, our findings demonstrate that CTCF binding to the wtAAV2 genome is necessary for proper viral gene expression and sufficient to enhance rAAV vector expression.

## RESULTS

### CTCF associates with wtAAV2 P5 and regulates viral gene expression

CTCF is known to regulate the life cycle of many DNA viruses (including herpesviruses, papillomaviruses and parvoviruses described above; [44–46, 50–52, 57, 61]). To determine whether CTCF binding sequences are present on the wtAAV genome, we performed *in silico* analysis on the wtAAV2, 5, and 8 genomes using the motif-scanning tool JASPAR (Fig. 1A, and Fig. S1; [62]). We found that all three wtAAV serotypes contained putative CTCF-binding sequences throughout their genomes. wtAAV2 only contained 3 predicted binding sites (Fig. 1A, red bars), while 9 were predicted for wtAAV8 (Fig. 1A, orange bars) and 7 were predicted for wtAAV5 (Fig. 1A, green bars). To broadly assess whether CTCF regulates wtAAV2 gene expression, we performed global knockdowns of CTCF using RNAi in HEK293T and U2OS cells, confirming the knockdown efficiency by western blot (Fig. 1B). Depletion of cellular CTCF levels by approximately 72% in HEK293T and 94% in U2OS cells resulted in a substantial decrease in *Rep68/78* transcripts during wtAAV2 mono-infection (Fig. 1C). Interestingly, the AAV2-CBE is conserved between wtAAV2 and wtAAV8 (Fig. 1A indicated by *). This positional conservation proximal to the P5 promoter (hereafter labeled as AAV2-CBE) led us to hypothesize that AAV2-CBE regulates Rep68/78 gene expression. We generated a mutant wtAAV2 lacking the CBE (wtAAV2^ΔCBE^; Fig. 1D) and quantified CTCF binding to the viral genome. ChIP-qPCR demonstrated that CTCF is bound to the predicted AAV2-CBE upstream of the P5 region in wtAAV2 and confirmed loss of binding at this site in wtAAV2^ΔCBE^ (Fig. 1E). Deletion of AAV2-CBE did not impact the production of wtAAV2 virus when HEK293T cells were co-transfected with the AdV helper plasmid (expressing E2A, E4orf6, and VA RNA) and an infectious clone plasmid (pIC) expressing wtAAV2 or wtAAV2^ΔCBE^ (Fig. 1F). However, loss of CTCF binding resulted in a decrease in *Rep68/78* transcripts during mono-infection of HEK293T and U2OS cells (Fig. 1G). These results showed that the CTCF binding at the AAV2-CBE is necessary for efficient viral gene expression.

**Figure 1:**
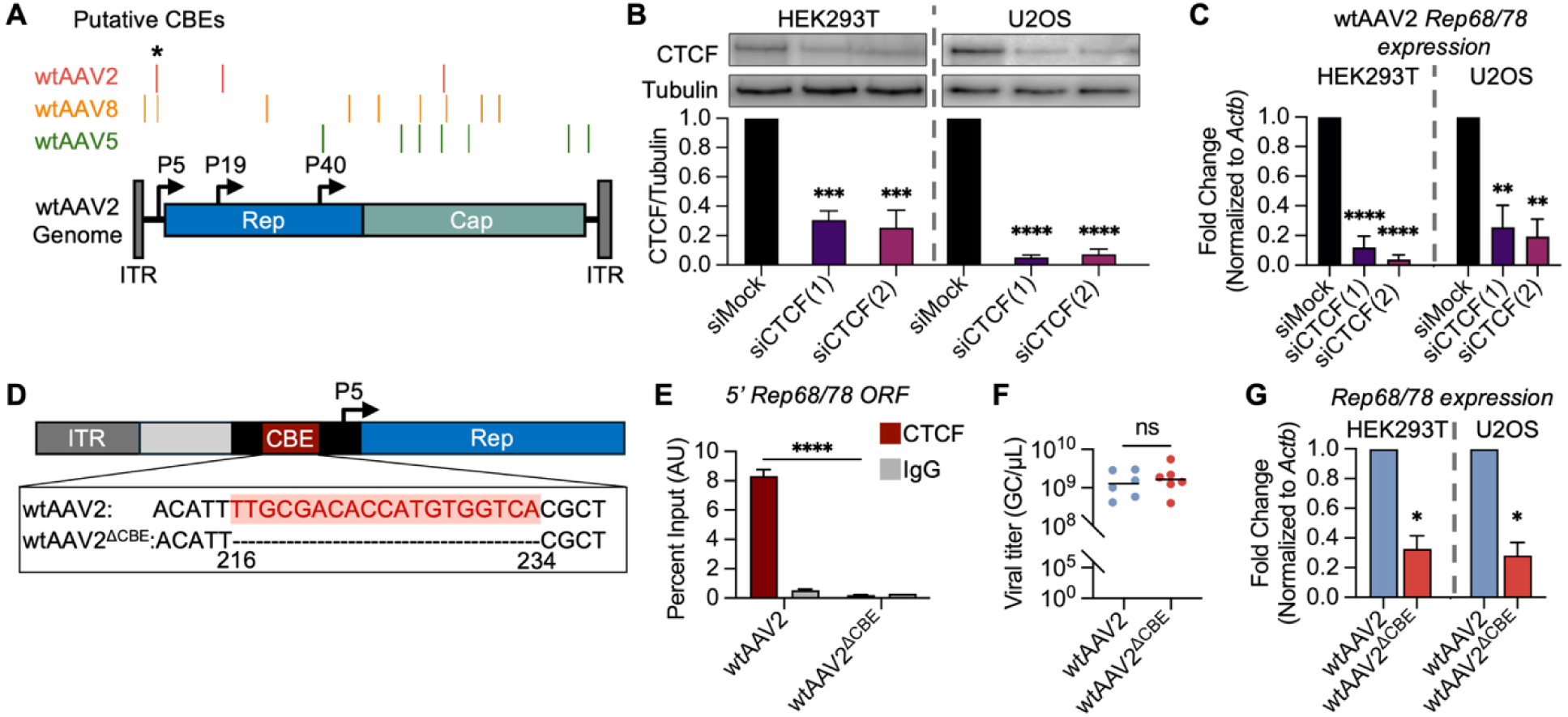
CTCF binding element regulates wtAAV2 gene expression. (A) *In silico* predicted CBEs on wtAAV2, 8, and 5 viral genomes. Approximately position is shown on the wtAAV2 genome schematic. * highlights conserved P5 CBE on wtAAV2 and wtAAV8. (B) HEK293T (left, n=4) and U2OS (right, n=3) cells treated with siRNA and infected with wtAAV2. Western blot analysis confirmed CTCF (top) knockdown by siRNA compared to cells transfected with a non-specific siRNA (siMock). Tubulin (bottom) was used as a loading control. Lower graph represents the quantification of CTCF levels relative to tubulin. (C) rt-qPCR analysis of Rep68/78 expression during siRNA treatment in HEK293T (left, n=4) and U2OS (right, n=3) cells. Expression fold change was calculated relative to cellular *Actb* levels. (D) Schematic showing the 5’ end of wtAAV2 genome and the AAV2-CBE mutated in wtAAV2^ΔCBE^ (red highlighted region indicates original sequence). Black box represents the P5 region with the arrow positioned at the TATA box. (E) ChIP-qPCR with primers complementary to the P5 promoter shows binding of CTCF (red) relative to non-specific IgG control antibody (grey). (F) wtAAV2 (blue) and wtAAV2^ΔCBE^ (red) titers represented as genome copies per µL (GC/µL). HEK293T cells co-transfected with IC and adenovirus helper plasmids were harvested at 72 hours and tittered using qPCR against the Rep68/78 gene. Individual dots represent a virus stock with black line shows the mean of each condition (n=6). (G) RT-qPCR analysis of Rep68/78 expression in wtAAV2 (blue) and wtAAV2^ΔCBE^ (red) in HEK293T (left, n=3) and U2OS (right, n=5) cells. All infections were carried out at an MOI of 5,000 vg/cell for 24 hours. Error bars represent standard error mean (SEM). Statistic *p*-value ns >0.5, * <0.03, ** <0.002, *** <0.0002, **** <0.0001.

### AAV2-CBE regulates wtAAV2 localization to host chromatin

Given CTCF’s role in host genome organization, regulation of chromatin landscape, and homologous recombination repair (HR; [63, 64]) and the importance of viral navigation of the host milieu for efficient expression and viral replication [65–67], we performed Viral Chromosome Conformation Capture (V3C-seq) to determine how CTCF might contribute to the localization of wtAAV2 genomes to the host genome[65, 68]. The V3C-seq assay captures the spatial association of viral genomes to host cellular sites using formaldehyde-mediated crosslinking of virus-host interactions. Subsequent high-throughput sequencing of unique virus-host hybrid DNA junctions generated by restriction enzyme digestion and intermolecular ligation enables comparison of wtAAV2 and wtAAV2^ΔCBE^ localization sites on the cellular genome. Analysis of the top 150 sites enriched for viral genomes revealed a wide distribution across all host chromosomes (Fig. 2A). Only 28.8% of wtAAV2 localization sites on the host were shared by wtAAV2^ΔCBE^, suggesting this CTCF binding element directs the ability of wtAAV2 genomes to navigate the nucleome (Fig. 2B, Jaccard test of statistical significance shown in Fig. 2C).

**Figure 2:**
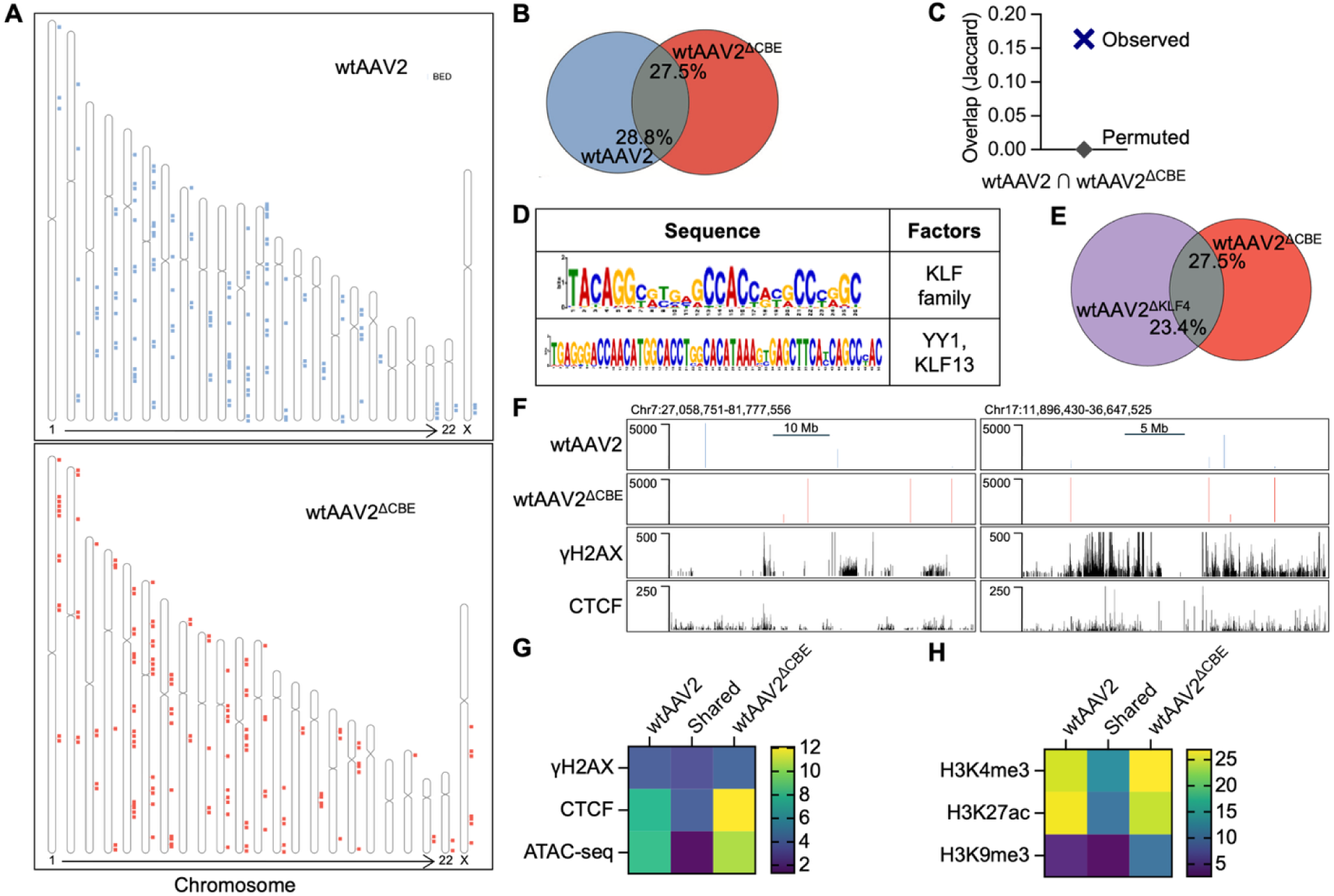
AAV2-CBE regulates wtAAV2 localization to host cellular sites. (A) Position of the top 150 virus-associated HindIII wtAAV2 (top) and wtAAV2^ΔCBE^ (bottom) enriched sites of localization from V3C-seq analysis represented as a dot across chromosomes (n=2). (B) Venn diagram of wtAAV2-associated (blue), wtAAV2^ΔCBE^-associated (red), and shared (wtAAV2 and wtAAV2^ΔCBE^, grey) localization sites on the host. Percentages represent the portion of wtAAV2 or wtAAV2^ΔCBE^ sites that are shared. (C) Jaccard values representing the extent of overlap compared to a “permuted” value of the intersection with randomly generated library of 150 fragments of 5kb size across the human genome. A value of 0 = no overlap, 1 = complete overlap. (D) Results of the *in silico* analysis using MEME and TOMTOM (ref) pipelines of the host sites of unique wtAAV2^ΔCBE^ localization. (E) Venn diagram of wtAAV2^ΔKLF4^ -associated (purple), wtAAV2^ΔCBE^-associated (red), and shared (wtAAV2^ΔKLF4^ and wtAAV2^ΔCBE^, grey) localization sites on the host. Percentages represent the portion of wtAAV2^ΔKLF4^ or wtAAV2^ΔCBE^ sites that are shared. (F) UCSC genome browser tracks showing representative localization of wtAAV2 and wtAAV2^ΔCBE^ on chromosome 1 (top) and chromosome 17 (bottom). Localization compared with UCSC tracks showing published γH2AX and CTCF ChIP-seq data. G,H) Heat map showing the percentage of wtAAV2-associated, wtAAV2^ΔCBE^-associated, or shared wtAAV2 and wtAAV2^ΔCBE^ - associated sites that colocalize with (G) γH2AX, CTCF ChIP-seq, and ATAC-seq peaks from published data sets and (H) H3K4me3, H3K27ac, and H3K9me3 ChIP-seq peaks from published ENCODE datasets.

Since wtAAV2^ΔCBE^ dramatically relocalizes to different sites on the host genome, we wanted to determine what distinguishes these sites. *In silico* analysis of consensus sequences enriched at sites of wtAAV2^ΔCBE^ localization showed binding sequences for the transcription factors YY1 and KLF protein family (Fig 2D). YY1 binds to and suppresses wtAAV2 P5, suggesting that viral relocalization to YY1-binding sites on the host may contribute to viral gene suppression [69, 70]. Notably, we recently showed that KLF4 binds to the P5 region of wtAAV2 and regulates viral localization to sites of DNA damage and viral gene expression[71]. This KLF4 site is positioned 69 bp upstream of the AAV2-CBE site in wtAAV2. Due to the proximity of the CBE and KLF4 site in the viral genome, we wanted to determine whether the loss of AAV2-CBE or the AAV2 KLF4-binding site directs the viral genome to associate with the same regions of the host genome. Intersection analysis showed that only 23.4% of wtAAV2^ΔKLF4^ sites were shared with wtAAV2^ΔCBE^ (Fig. 2E). This suggested that wtAAV2 CTCF-directed localization is distinct from KLF4-mediated localization.

Previous work has established that wtAAV2 localizes to sites of host DNA damage marked by γH2AX and that this localization aids viral gene expression [66, 71]. To determine if the decrease in Rep68/78 expression is associated with an attenuated localization to DNA damage sites, we compared the percentage of wtAAV2 unique, wtAAV2^ΔCBE^ unique, and shared sites that overlapped with γH2AX ChIP-seq peaks. While loss of CTCF binding resulted in overall relocalization of wtAAV2, it did not affect the frequency of localization to sites of DNA damage (visualization with UCSC genome tracks, Fig. 2F, and quantification 2G top row). We recently showed an enrichment of *in silico* predicted CTCF binding sites at wtAAV2-associated genomic regions [71]. To determine if loss of AAV2-CBE precluded localization to CTCF sites on the host genome, we assessed the colocalization of wtAAV2 with published CTCF ChIP-seq peaks. Interestingly, we observed an increase in colocalization to CTCF sites on the host, with only 8% of unique sites associated with wtAAV2 compared with 12% of wtAAV2^ΔCBE^- associated sites. However, only ∼4% of the sites that retained wtAAV2 localization upon AAV2-CBE deletion intersected with CTCF ChIP-seq sites (Fig. 2G middle row). These observations suggested that additional host factors or sequence motifs contribute to wtAAV2 localization to cellular CTCF sites.

As a multifunctional protein, CTCF aids in chromatin accessibility and the establishment of active chromatin marks on the cellular and viral genomes[31, 72, 73]. We therefore assessed how wtAAV2/wtAAV2^ΔCBE^ localizes to the host genome relative to accessible chromatin regions. Using publicly available ATAC-seq data, we found that 8% of unique localization sites of wtAAV2 and 10% of wtAAV2^ΔCBE^ unique localization sites overlapped with open chromatin regions in the host. However, only 1% of these localization sites shared by both viruses correlated with open chromatin regions on the host (Fig. 2G bottom row). This suggested that while wtAAV2 can localize to open chromatin on the host genome independent of the CBE, the lack of shared open chromatin regions suggested an additional layer driving colocalization.

To investigate this further, we assessed the colocalization of specific histone marks and wtAAV2/wtAAV2^ΔCBE^ association sites. Cellular regions shared by both wtAAV2 and wtAAV2^ΔCBE^ localization had a similarly high enrichment (average of 25% each) of histone marks associated with transcriptionally active chromatin (H3K4me3 and H3K27ac; Fig. 2H top and middle rows). However, histone marks associated with transcriptionally repressive chromatin, H3K9me3, colocalized with 10% of wtAAV2^ΔCBE^ unique sites compared to only 4% of wtAAV2 unique sites and 2% of shared sites (Fig. 2H bottom row). These observations indicated that while the presence of the CBE is not sufficient to ensure association with active chromatin domains, it may still play a role in shielding the wtAAV2 genome from associating with repressive chromatin environments.

### CBE deletion alters wtAAV2 chromatin landscape

Since wtAAV2-CBE mediates viral localization and interactions with different chromatin domains on the host genome, we hypothesized that this element regulates the viral chromatin landscape. Indeed, in large DNA viruses like HSV-1 and EBV, CTCF binding restricts the spread of inactive chromatin marks to regulate viral gene expression[46, 72, 73]. To determine if CTCF similarly regulates wtAAV2 expression through alterations to the viral chromatin environment, we performed CUT&RUN analysis, measuring the changes in active (H3K4me3 and H3K27ac) and inactive (H3K9me3) histone marks (Fig. 3A). Quantification of the mean IgG normalized RPM (reads per million) across the genome between the ITRs showed a decrease in both active (H3K4me3 and H3K27ac) and repressive (H3K9me3) marks in wtAAV2^ΔCBE^ (Fig. 3B). However, when we assessed the changes in chromatin marks around the P5 promoter, we observed a 13-fold decrease in H3K4me3, and 3-fold decrease in H3K27ac, and a doubling of H3K9me3 when the AAV2-CBE was deleted (Fig. 3C). This suggested that CTCF binding at AAV2-CBE impacts both the local and global chromatin. Although lower, the continued presence of active chromatin marks in wtAAV2^ΔCBE^ may explain why viral gene expression was not abolished upon loss of AAV2-CBE (observed in Fig. 1G). Additionally, the limited expression of wtAAV2 gene expression during mono-infection may account for the observation of H3K9me3 across the viral genome.

**Figure 3:**
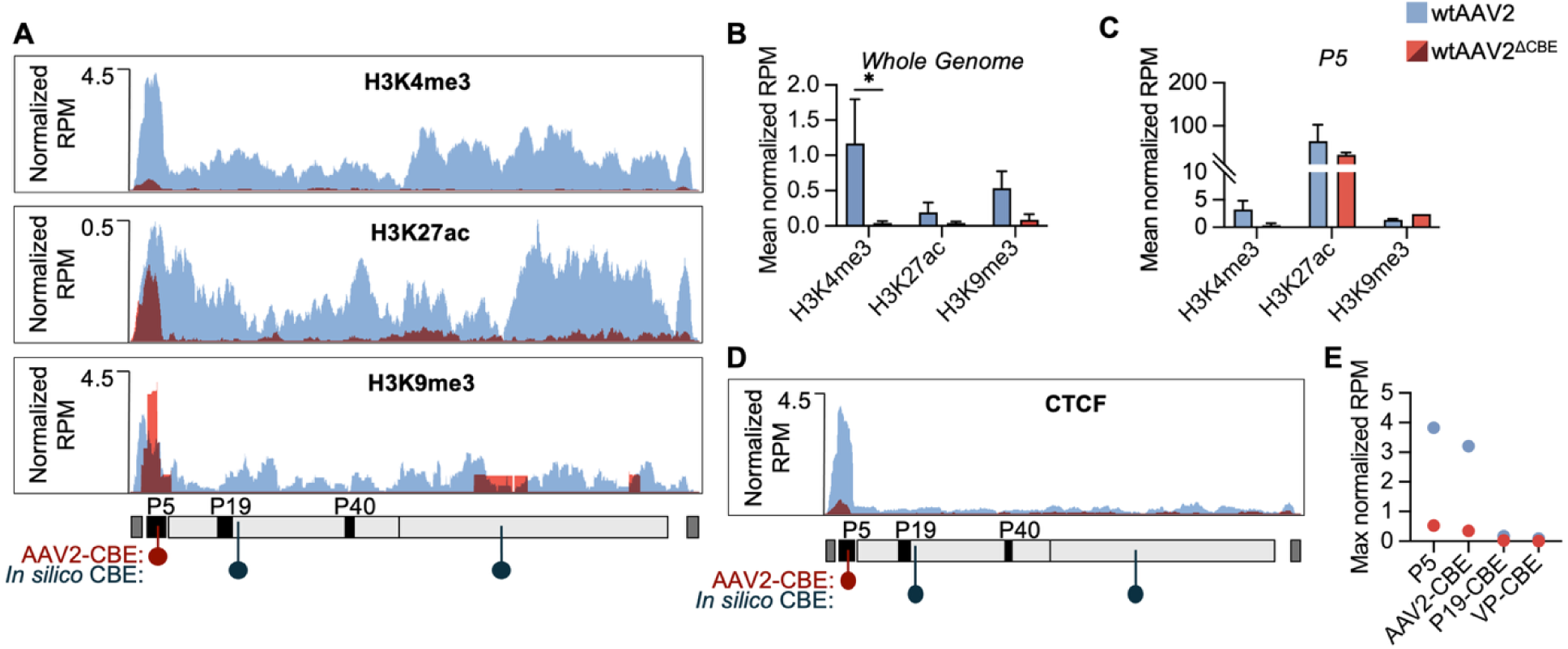
AAV2-CBE influences wtAAV2 histone modifications. (A-E) wtAAV2 (blue) and wtAAV2^ΔCBE^ (red) CUT&RUN profiling of HEK293T cells infected at 500 MOI for 24 hours. (A) wtAAV2 and wtAAV2^ΔCBE^ (dark red indicates overlapped regions) profiles are overlayed on a single UCSC genome browser track showing IgG normalized reads per million (RPM) for H3K4me3, H3K27ac, and H3K9me3. Tracks are plotted on the wtAAV2 genome (bottom schematic) showing ITRs (dark grey), REP and CAP regions (light grey), promoter regions (black box), in silico predicted CBEs (dark blue popsicles), and wtAAV2 AAV2-CBE location (red popsicle). (B) Quantification of the mean normalized RPM across the genome minus the ITRs for each histone mark. (C) Quantification of the mean normalized RPM in the P5 region (black box on schematic in A). (D) UCSC genome browser track of CUT&RUN CTCF binding along the wtAAV2 genome. (E) Comparison of the maximum height of normalized RPM peak in the region indicated. wtAAV2 n=2, wtAAV2^ΔCBE^ n=3, wtAAV2^ΔCBE^ H3K9me3 n=1. Error bars represent standard error mean (SEM). Statistic *p*-value ns (not shown) >0.5, * <0.03.

Since we observed that wtAAV2 contains three predicted CTCF-binding sequences across the genome (Fig. 1A), we experimentally assessed CTCF binding using CUT&RUN analysis. While we observed a strong CTCF-binding peak at the AAV2-CBE at the P5 promoter in wtAAV2, we did not detect CTCF binding at the *in silico* predicted P19 or VP CBEs (Fig. 3D, 3E). In wtAAV2^ΔCBE^, we observed a 9-fold decrease in CTCF binding at AAV2-CBE. However, this was not associated with a change in CTCF binding to the other predicted CBEs in wtAAV2. These findings suggested that, unlike other viral systems, wtAAV2 contains a single CTCF-bound CBE that is necessary to regulate wtAAV2 gene expression and chromatinization during mono-infection.

### AAV2-CBE enhances transgene expression from rAAV vectors

rAAV vectors are engineered to retain only the AAV-ITRs, whereas the cis-regulatory elements between the ITRs (including the AAV2-CBE) are excluded from current rAAV modalities [1, 3, 74]. To determine if the AAV2-CBE is sufficient to regulate transgene expression, we engineered rAAV2 vectors expressing a GFP cassette driven by the CMV promoter (CMV-eGFP, rAAV2) to contain the AAV2-CBE sequence (rAAV2^CBE^ Fig. 4A). Binding of CTCF to this site was confirmed by ChIP-qPCR (Fig. 4B). Addition of the AAV2-CBE in rAAV2 did not impact vector production in HEK293T cells co-transfected with plasmids expressing AdV helper, wtAAV2 Rep-Cap, and the pIC of rAAV2 or rAAV2^CBE^ (Fig. 4C). To determine the impact of AAV2-CBE on rAAV2 transgene expression, we transduced HEK293T cells with rAAV2 and rAAV2^CBE^ at increasing MOIs (10, 500, and 5000 vg/cell) and assessed GFP expression by western blot. While GFP expression was undetectable until 500 MOI in rAAV2-transduced cells, low levels of expression were seen at 10 MOI in rAAV2^CBE^– transduced cells (Fig. 4D, lanes 2 vs 5). At high MOIs however, similar levels of GFP expression were seen between rAAV2 and rAAV2^CBE^ (Fig. 4D, lanes 4 vs 7). Transduction with rAAV2 or rAAV2^CBE^ did not impact global CTCF levels (Fig. 4D).

**Figure 4:**
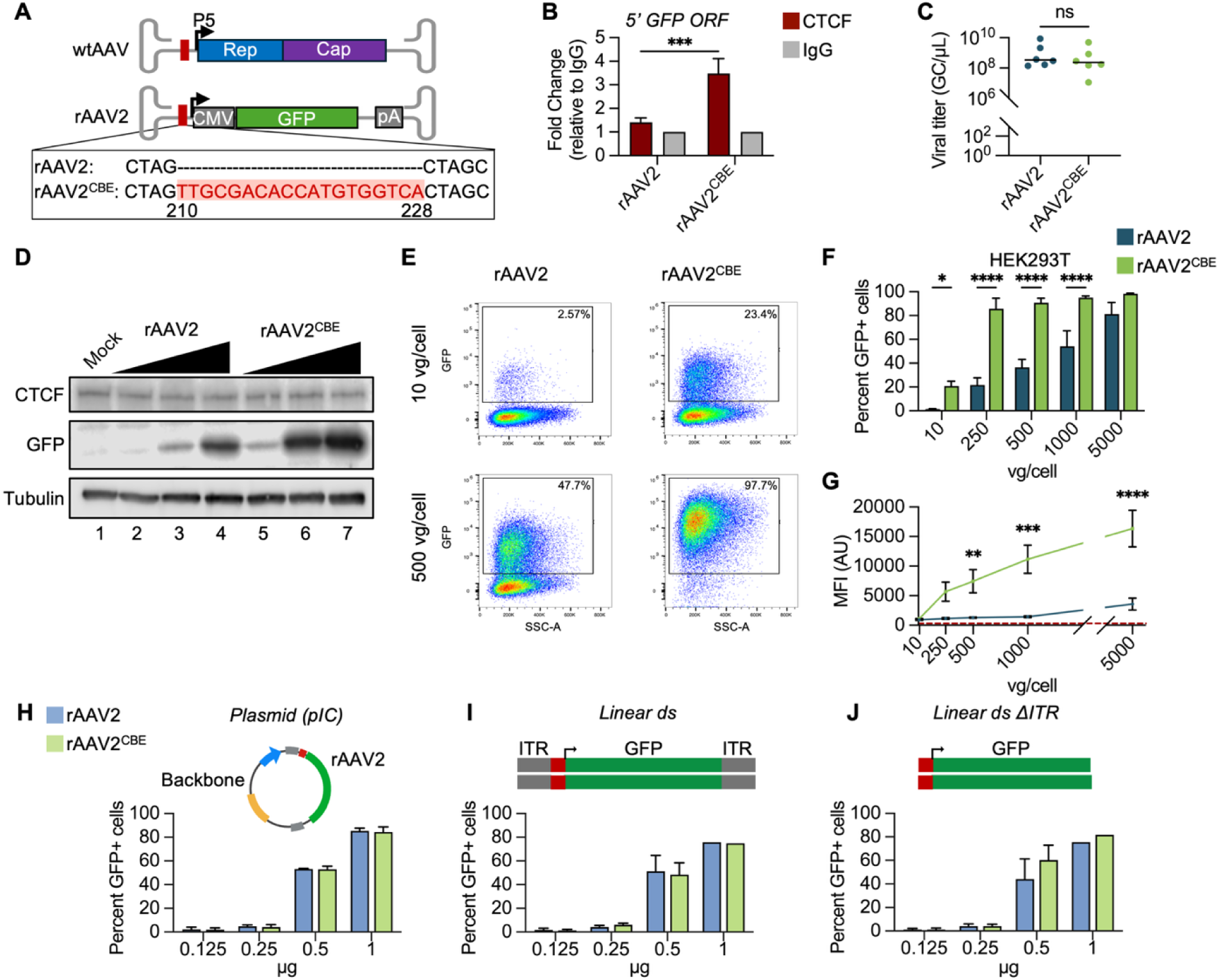
CTCF binding elements are sufficient to enhance rAAV2 transgene expression in 293T cells. (A) Schematic comparing the wtAAV2 and rAAV2. Red box and highlight indicate sequence and position of AAV2-CBE in rAAV2. (B) ChIP-qPCR fold change analysis of CTCF (red) relative to IgG control (grey) with primers complementary to the CMV promoter. HEK293T cells transduced for 24 hours at 5,000 MOI (n=10). (C) rAAV2 (teal) and rAAV2^CBE^ (green) titers represented as genome copies per µL (GC/µL). HEK293T cells co-transfected with IC, adenovirus helper, and Rep and Cap expressing plasmids were harvested at 72 hours and tittered using qPCR against the GFP gene. Individual dots represent a virus stock with black line shows the mean of each condition (n=6). (D) Western blot analysis of HEK239T cells 24 hours post rAAV2, rAAV2^CBE^, or mock transduction at 10, 500, and 5,000 genome copies/cell (GC/cells). Levels of CTCF (top) and GFP (middle) are shown with tubulin (bottom) as a loading control. (E) Representative flow cytometry plots for GFP-A against SSC-A for rAAV2 and rAAV2^CBE^ in HEK293T cells transduced for 24 hours at 10 and 500 GC/cell. Gating for GFP positive cells based on mock samples shown by black box with the percentage of single cells expressing GFP in the top right. (F) Quantification of flow cytometry percentage of GFP positive single cells over increasing MOI comparing rAAV2 and rAAV2^CBE^ (n≥4). (G) Median Fluorescent Intensity of GFP in arbitrary units from flow cytometry over increasing MOI. (n≥4). Red dashed line represents average mock MFI. (H-J) Flow cytometry of HEK293T cells 24 hours after transfection with increasing µg of DNA in the form pIC (H), double stranded (ds) linear genome with (I) and without (J) the ITRs. No statistical difference was seen between rAAV2 (light blue) and rAAV2^CTCF^ (green) in any condition (n=2 except 1µg n=1). Error bars represent standard error mean (SEM). Statistic *p*-value ns/not shown >0.5, * <0.03, ** <0.002, *** <0.0002, **** <0.0001.

To investigate the transduction efficiency of rAAV2^CBE^ at the single-cell level, we performed flow cytometry (Fig. 4E-4G). The addition of the AAV2-CBE in rAAV2 dramatically increased the percentage of HEK293T cells expressing the GFP transgene. At 10 MOI, where GFP was almost undetectable in rAAV2, rAAV2^CBE^ had an 18-fold increase in transduction (Fig. 4F and Fig. S2). By 250 MOI, rAAV2^CBE^ had reached near saturation with 85% of cells expressing GFP. By comparison, rAAV2 had significantly lower transduction efficiencies till they reached a transduction MOI of 5,000 vg/cell, where a no difference with rAAV2^CBE^ transduced cells was detectable. Using median fluorescent intensity (MFI) to assess transgene expression, we observed that, in addition to transduction efficiency, rAAV2^CBE^ vectors had higher overall levels of expression compared to rAAV2 (Fig. 4G). Interestingly, at 5,000 MOI, when both vectors showed similar percentages of GFP-positive cells, rAAV2^CBE^ had a significantly higher MFI. Taken together, the western blot and flow cytometry analyses suggested that the addition of AAV2-CBE in rAAV2 is sufficient to increase both vector transduction and expression in 293T cells.

To determine whether CTCF-mediated enhancement of transgene expression can be generalized to heterologous DNA molecules, we transfected HEK293T cells with increasing amounts of circularized rAAV2 or rAAV2^CBE^ pIC. Flow cytometry analysis showed no significant difference between GFP expression in prAAV2- and prAAV2^CBE^ (Fig. 4H). To determine whether this CBE impacts the expression of transfected linear viral genomes, we isolated the double-stranded viral genome from pIC by restriction enzyme digestion. As shown in Fig. 4I and 4J, addition of AAV2-CBE did not impact GFP expression from double-stranded linear rAAV2 when it contained (Fig. 4I) or lacked (Fig. 4J) the ITR. These observations suggested that the impact of AAV2-CBE on transgene expression is specific to capsid-delivered rAAV2 genomes.

### Generalizability of CBE-mediated enhancement of rAAV-delivered transgene expression

To assess if CTCF-mediated enhancement of transgene expression is generalizable to additional tissue types, we performed these studies in U2OS and HepG2 cells. Since HEK293T cells constitutively express the AdV helper proteins E1A/B, we wanted to determine whether these viral proteins are impacting the increase in rAAV2^CBE^ transgene expression (observed in Fig. 4). Transduction of U2OS cells demonstrated a phenotype like that of HEK293T cells, with rAAV2^CBE^ vectors yielding significantly higher percentage of GFP positive cells (Fig. 5A). While overall MFI in U2OS was lower than in HEK293T cells, GFP expression was significantly higher in AAV2^CBE^ compared to rAAV2 at high MOIs (Fig. 5B). We additionally tested vector transductions in the liver cell line HepG2, representative of liver tissue, which is a key target in multiple rAAV gene therapy applications [75]. Although HepG2 cells demonstrated lower overall transduction efficiency with rAAV2 vectors, addition of the AAV2-CBE increased transduction and GFP expression (Fig. 5C, 5D, respectively). To determine whether CTCF-mediated enhancement of transgene expression extends to therapeutically relevant model systems, we transduced induced pluripotent stem cell-derived retinal pigment epithelium (iPSC-RPE) cells for 48 hours. rAAV2^CBE^ transduced iPSC-RPE cells on average 1.45-fold higher than rAAV2 across increasing MOIs (Fig. 5E). Additionally, rAAV2^CBE^ had higher GFP expression in iPSC-RPEs, as seen by comparison of MFI (Fig. 5F). These transduction studies across immortalized and iPSC-derived cells demonstrate that the AAV2-CBE is sufficient to enhance transgene expression in multiple cellular contexts.

**Figure 5:**
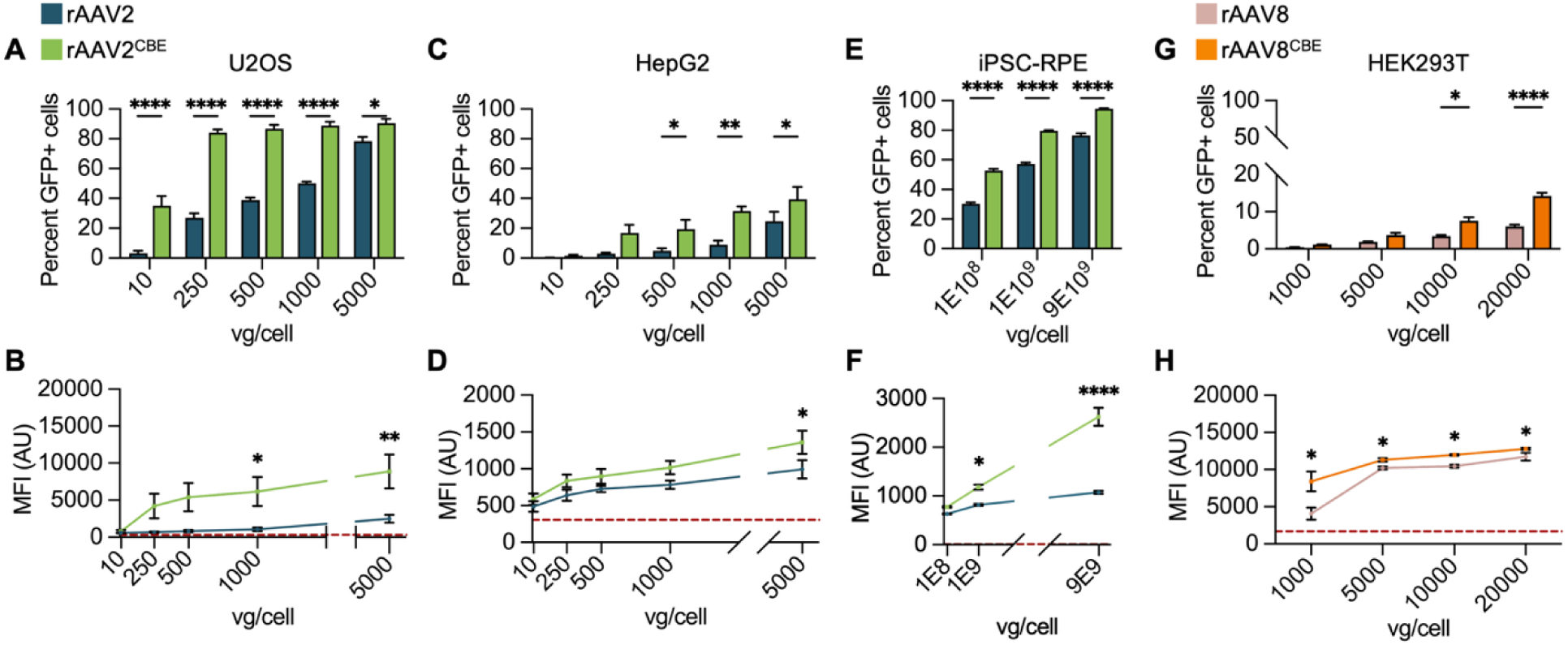
CTCF binding elements are sufficient to enhance transgene expression in independent cell types and when delivered with AAV8 capsid. Flow cytometry analysis of cells transduced with indicated increasing MOI 24hpt. (A,C,E,G) Quantification of flow cytometry percent of single cells expressing GFP. (B,D,F,H) Median Fluorescent Intensity of GFP in arbitrary units from flow cytometry over increasing MOI. (n≥3). Red dashed line represents average mock MFI. (A,B) U2OS cells transduced with rAAV2 (teal) and rAAV2^CBE^ (green). (C,D) HepG2 cells transduced with rAAV2 and rAAV2^CBE^. (E,F) Retinal pigment epithelial iPSC-derived cells transduced with rAAV2 (teal) and rAAV2^CBE^ (green). (G,H) HEK293T cells transduced with rAAV8 (pink) and rAAV8^CBE^ (orange). Error bars represent standard error mean (SEM). Statistic *p*-value ns >0.5, * <0.03, ** <0.002, *** <0.0002, **** <0.0001.

Clinical applications of rAAV gene therapy rely on the delivery of the transgene using multiple natural capsid types and newly engineered capsids that possess distinct tissue-specific tropism [76]. Additionally, the AAV capsid has been shown to mediate epigenetic control of rAAV gene expression[77]. Therefore, we assessed the impact of the AAV2-CBE on transgene expression when delivered using a clinically relevant capsid. We produced rAAV vectors (as described above) packaged into the AAV8 capsid (labeled rAAV8 and rAAV8^CBE,^ respectively) and assessed GFP expression by flow cytometry. While the transduction efficiency of rAAV8 was significantly lower than that of rAAV2 in HEK293T (Fig. S3A), rAAV8^CBE^ had significantly higher transduction efficiency at 10,000 and 20,000 MOI compared to rAAV8 (Fig. 5G). Even at the lower MOIs tested (1,000 and 5,000 MOI), rAAV8^CBE^ transduction lead to a higher MFI compared to rAAV8 (Fig. 5H). We additionally tested the AAV9 capsid and found that, while there was a slight increase in transduction in HEK293T cells at high MOIs, there was no significant difference in percentage of GFP-positive cells or in MFI between rAAV9 and rAAV9^CBE^ (Fig. S3B and S3C). Overall low transduction efficiency may be due to the poor tropism of AAV8 and AAV9 in HEK293T cells[78]. Despite these observations, our findings suggested that the impact of AAV2-CBE on rAAV gene expression is at least partially independent of the AAV capsid.

### rAAV2 chromatin landscape is altered by the addition of AAV-CBE

Despite being designed to retain only the wtAAV2 ITRs, *in silico* analysis revealed the rAAV2 genome harbors one predicted CBE in the CMV enhancer and five in the 3’ GFP and polyA region. CUT&RUN analysis confirmed binding of CTCF at the engineered CBE site in rAAV2^CBE^ (Fig. 6A). rAAV2^CBE^ had an increased max RPM peak (Fig. 6B) and mean RPM (Fig. 6C) over the 19bp AAV2-CBE region compared to rAAV2, corroborating our earlier ChIP-qPCR assays (Fig. 4B). Notably, the CTCF CUT&RUN analysis revealed dynamic CTCF binding throughout the rAAV2 and rAAV2^CBE^ genomes.

**Figure 6:**
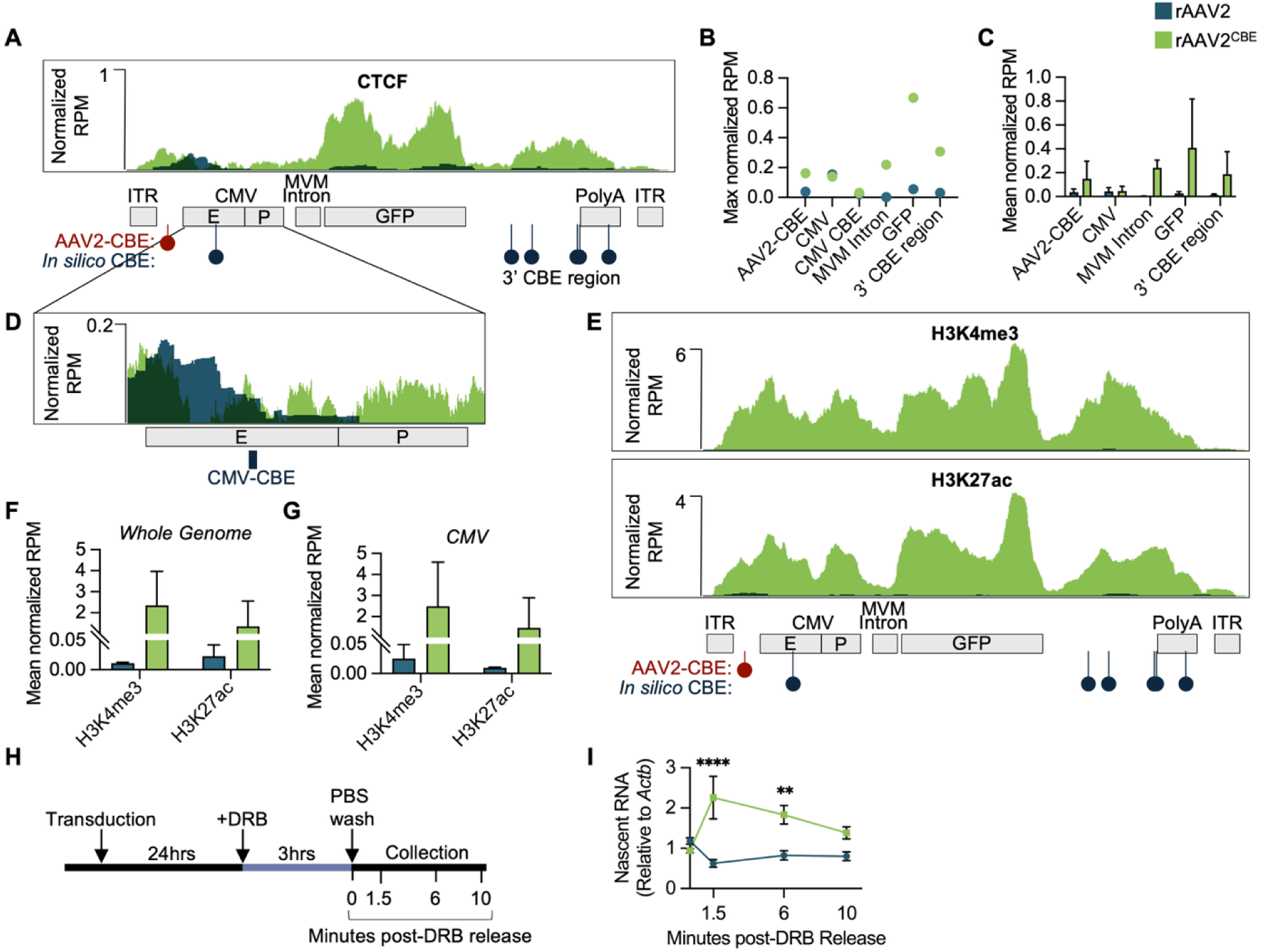
AAV2-CBE regulates rAAV2 chromatin landscape and enhances nascent RNA production. (A-E) rAAV2 (teal) and rAAV2^CBE^ (green) CUT&RUN profiling of HEK293T cells transduced at 500 MOI for 24 hours. (A) rAAV2 and rAAV2^CBE^ profiles are overlayed on a single UCSC genome browser track showing IgG normalized reads per million (RPM) for CTCF binding. Tracks are plotted on the rAAV2 genome (bottom schematic) with in silico predicted CBEs (dark blue popsicles), and inserted AAV2-CBE location (red popsicle). (B) Comparison of the maximum height of normalized RPM peak in the region indicated. (C) Quantification of the mean normalized RPM for each location indicated. (D) Inset of the CMV enhancer-promoter region of rAAV2 and rAAV2^CBE^ CUT&RUN UCSC browser tracks. Black box is the region of CMV-CBE. (E) rAAV2 and rAAV2^CBE^ profiles are overlayed on a single UCSC genome browser track showing IgG normalized reads per million (RPM) for H3K4me3 and H3K27ac. (F) Quantification of the mean normalized RPM across the genome minus the ITRs for each histone mark. (G) Quantification of the mean normalized RPM in the CMV enhancer/promoter region. rAAV2 n=2, rAAV2^CBE^ H3K4me3 n=3, wtAAV2^CBE^ H3K27ac n=2. (H) Schematic of the timeline of DRB treatment and rAAV2 transductions. HEK293T cells were transduced at 1,000 GC/cell with rAAV2 or rAAV2^CBE^ for 24 hours. DRB was added at 40g/ml for 3 hours before cells were released with PBS wash and incubated in fresh medium for indicated time points. (I) rt-qPCR analysis of nascent RNA transcripts at indicated time points (n=4). Error bars represent standard error mean (SEM). Statistic *p*-value ns (not shown) >0.5, * <0.03, ** <0.002, *** <0.0002, **** <0.0001.

Interrogation of the CMV promoter region and the predicted CMV-CBE revealed a broad CTCF-binding peak at the 5’ end of the promoter with the 3’ end encompassing the CMV-CBE in rAAV2 (Fig. 6D, dark blue histogram). However, in rAAV2^CBE^, CTCF binding is shifted rightwards toward the AAV2-CBE, generating a sharper CTCF-binding peak at the CMV-CBE and a broad CTCF-binding peak over the upstream promoter sequence (Fig. 6D, green histogram). While there is no difference between the maximum peak height and mean coverage in these areas, the sliding of CTCF binding suggested the possibility of competitive recruitment (Fig. 6B and C).

The 3’ region of the viral genome, which contains five putative CBEs, demonstrated an enhancement of CTCF binding in rAAV2^CBE^ upon insertion of the AAV2-CBE to the 5’ region of the genome compared to rAAV2 (Fig. 6A). In rAAV2^CBE^, binding to the 3’ CBE region had a higher maximum peak height than at the CBE insert (Fig. 6B, 6C). The overall increase of CTCF binding to the predicted CBEs in the presence of the engineered AAV2-CBE suggests the potential of an overall cooperative effect on the rAAV2^CBE^ genome.

rAAV2 vectors typically contain the MVM viral capsid intron-1 (VP1) which contains a CTCF binding sequence[79]. In MVM, CTCF binding is required for proper processing of viral RNAs and in rAAV the addition of this intronic region increases transgene expression[57, 79]. Despite the presence of CTCF binding in MVM, our *in silico* and CUT&RUN analysis did not detect a CBE or CTCF binding to the MVM intron in rAAV2 (Fig. 6A). However, CTCF binding was detected at the MVM intron in rAAV2^CBE^ (Fig. 6A-6C), suggesting that this binding is dependent on the presence of the engineered AAV2-CBE.

Outside of the known and predicted CBEs, CUT&RUN analysis revealed low levels of CTCF binding in the GFP-coding region of rAAV2 (Fig. 6A-C). Binding in these regions was strongly enriched in rAAV2^CBE^, further strengthening our observation of an overall cooperative effect of incorporating AAV2-CBE into rAAV2. Additionally, this highlights the need to screen for CBEs inherent to transgenes, as they might regulate vector transgene expression.

We sought to determine if CTCF was sufficient to alter the rAAV2 vector chromatin landscape, thereby regulating gene expression. We performed CUT&RUN on rAAV2 and rAAV2^CBE^ vector genomes in HEK293T cells for active histone marks H3K4me3 and H3K27ac. Like its impact on wtAAV2, we found that the addition of the AAV2-CBE resulted in global changes to the viral chromatin environment (Fig. 6E). Interestingly, the addition of CTCF binding in rAAV2 caused a global increase in the mean RPM across the genome and specifically at the CMV promoter (Fig. 6F, 6G). The finding suggested CTCF is sufficient to regulate the chromatin environment of rAAV.

To interrogate if AAV2-CBE is increasing expression through facilitating RNA PolII activity, we compared nascent RNA production of the rAAV2 and rAAV2^CBE^ vectors. HEK293T cells were transduced at 500 MOI for 24 hours before transcriptional elongation was paused by treatment with 5,6-dichlorobenzimidazole 1-β-d-ribofuranoside (DRB, Fig. 6H). Immediately upon release, nascent RNA production was significantly higher in rAAV2^CBE^ compared to rAAV2 (Fig. 6I; 1.5 minutes post release). While nascent RNA production in rAAV2^CBE^ gradually declined over time, levels remained higher in rAAV2^CBE^– transduced cells than rAAV2 (Fig. 6I). Taken together, these results suggest that CTCF drives increased gene expression by establishing active chromatin marks and increasing transcription rates.

## DISCUSSION

In this study, we have demonstrated that CTCF-binding elements are both necessary for proper wtAAV2 gene expression and sufficient to confer higher expression potential in rAAV vectors. These changes in viral (or vector) gene expression are attributed to predominantly higher levels of active chromatin. The absence of AAV2-CBE from the wtAAV2 genome alters its ability to localize to repressive chromatin compartments, possibly facilitated by the “like-interacts-with-like” phenomenon of chromatin interactions [80, 81]. Taken together, our findings reveal new ways to enhance rAAV vector efficiency by modulating their chromatin landscape.

Understanding how rAAV is chromatinized has garnered recent interest as this can be used to improve gene therapy vector platforms. Identification of the NP220/HUSH complex establishing repressive chromatin marks and capsid-mediated control of viral histone mark deposition in a species-dependent manner have demonstrated that the wtAAV2 genome is chromatinized and contains modifiable histone subunits[77, 82, 83]. However, these studies used rAAV vectors containing exogenous promoters, delivered via modified capsid serotypes. Using the native wtAAV2 virus, our identification of CTCF-mediated regulation of viral gene expression opens the door to modulating rAAV gene expression by engineering the vector chromatin.

Recent studies have raised the possibility of the viral capsid regulating the lifecycle of wtAAV2 by controlling gene expression. In this regard, “dead zone” mutations in the capsid structure inhibit second-strand synthesis and subsequent viral transcription[84]. Interestingly, species-specific capsids dictate host range by modifying the histones associated with AAV genomes. However, our findings that the AAV2-CBE enhanced transgene expression when delivered using the wtAAV8 capsid suggested that the impact of CTCF may be independent of the viral capsid. However, conflicting results in rAAV9 suggest a potential interplay between capsid and CTCF-driven chromatin control. The contribution of genomic elements versus structural proteins to rAAV chromatin landscape requires further mechanistic dissection.

One of the most striking facets of our engineered CTCF-containing rAAV vectors is their effectiveness at low doses over conventional rAAV vectors. This application of a *cis*-regulatory element from wtAAV2 to enhance rAAV vector expression builds on our recent findings that the addition of a KLF4 binding site from wtAAV2 also enhanced rAAV vector transductions[71]. Interestingly, both KLF4 and CTCF binding elements are in the P5 region of the wtAAV2 genome, which also includes binding sites for other regulatory proteins (such as Rep and YY1) that control viral gene expression [35]. Our findings further highlight the complex interplay between viral and host factors in regulating wtAAV2 expression. Importantly, while KLF4 binding drives genome localization to cellular DDR sites, CTCF binding determines the viral chromatin landscape. However, the relative closeness of these sequences within the P5 region suggests a potential interplay between these factors. KLF4-mediated regulation of wtAAV2 is dependent on its interaction with the DNA damage protein PARP1[71]. CTCF function can be regulated in part through post-translational modifications (including PARylation) by PARP1, as observed in EBV [46, 61]. The recent findings of KLF4-PARP1 regulation of wtAAV2 opens the door to an additional layer of control. However, the lack of KLF4 binding sites in rAAV2 suggests that CTCF can act independently of KLF4 to govern the viral chromatin landscape. Future cooperative studies are necessary to understand the CTCF-KLF4-PARP1 relationship as a method to further enhance rAAV transgene expression. These elements add to the growing list of *cis*-acting sequences present in AAV but not included in current rAAV vectors, which can enhance vector-mediated transgene expression.

In host and DNA viral genomes, CTCF is known to regulate gene expression by controlling 3D chromatin structure through the formation of promoter-enhancer loops [31]. Previous studies have identified the potential of a short-range genomic loop in wtAAV2 between Rep68-bound P5 and Sp1-bound P19[35]. However, these structures were observed in vitro with a wtAAV2 DNA substrate incubated with purified recombinant proteins. The question remains whether wtAAV and rAAV genomes can be packaged into histones and form short chromatin loops that can overcome steric hindrance. Due to the high abundance of viral and vector genomes used in laboratory settings, it remains technically challenging to determine which viral genomes are chromatinized, which are looped, and which serve as platforms for active transcription. Canonically, CTCF drives the formation of genome loops via the loop extrusion model [38, 40]. According to this model of looping, distal DNA elements are brought together by homodimerization of CTCF molecules, stabilized by Cohesin [39]. While our CTCF CUT&RUN analysis shows a single peak of CTCF binding on wtAAV2, this does not preclude the possibility of a chromatin loop, either facilitated through interaction with other transcription factors (as seen with a CTCF-YY1 loop in HPV[51]) or dimerization between multiple CTCF-bound sites in concatemerized wtAAV2 episomes.

While our CTCF CUT&RUN analysis shows a single peak of CTCF binding on wtAAV2, this does not preclude the possibility of a chromatin loop either facilitated through interaction with other transcription factors (as seen with a CTCF–YY1 loop in HPV [51]) or dimerization between multiple CTCF-bound sites in concatemerized wtAAV2 episomes. Notably, CTCF’s position adjacent to the ITR and its established role in homologous recombination raise the possibility that CTCF may also regulate concatemer formation itself through the ITRs [63, 64, 85], adding a second mode by which CTCF could influence the extrachromosomal fate of wtAAV2 genomes. Interestingly, we showed that our rAAV2-GFP construct contained additional sites of CTCF binding across the genome, which were enhanced by the addition of the AAV2-CBE. Additionally, the Cohesin subunit SMC1 has been found to associate with Rep and viral replication centers[86], suggesting these architectural protein complexes might synergize with CTCF, presenting an enticing model for regulating parvovirus gene expression by 3D looping. Taken together, our identification of CTCF as a regulator of wtAAV2 gene expression builds on the existing model of DNA viruses in utilizing key host proteins. We further demonstrate how this can be leveraged to enhance rAAV gene expression with the potential to advance current gene therapy strategies.

## MATERIALS AND METHODS

### Cell lines

Human embryonic kidney cells (HEK293T) and female human U2OS cells in Dulbecco’s modified Eagle’s medium (DMEM, high glucose; Gibco) supplemented with 5% Serum Plus (Sigma Aldrich) and 50 µg/ml gentamicin (Gibco). Human hepatocyte HepG2 cells were maintained on plates seeded with 50 mg/mL rat tail collagen (BD Biosciences) in 0.02 N acetic acid solution in Dulbecco’s modified Eagle’s medium/nutrient mixture F-12 (DMEM/F-12, high glucose; Gibco) supplemented with 5% Serum Plus (Sigma Aldrich) and 50 μg/mL penicillin/streptomycin (Gibco). All cells were cultured at 37°C and 5% CO_2_.

### iPSC-RPE differentiation

Induced pluripotent stem cell-derived retinal pigment epithelium (iPSC-RPE) cells were differentiated as described previously [87] from an iPSC line with homozygous mutation in *BEST1* (c.598C>T; p.R200X). Briefly, at the start of differentiation (day 0), ReLeSR was used to lift iPSCs and form aggregates called embryoid bodies (EBs). Over the first four days, the suspension culture of EBs, initially prepared in mTeSR Plus, were gradually transitioned to Neural Induction Medium (NIM; 500 mL DMEM/F12 (1:1), 1% N2 supplement, 1% MEM non-essential amino acids, 1% L-glutamine, 2 mg/mL heparin). On day 7, Laminin (Cat# 23017015; 1:20 dilution prepared in DMEM/F12) coated 6-well culture plates were used to plate EBs in the NIM media and media was changed regularly. On day 16, the 3D neurospheres were mechanically lifted off and the cells that remained adhered were maintained in the retinal differentiation media (RDM; DMEM:F12 (3.5:1.5), 2% B27 without retinoic acid, 1% antibiotic-antimycotic solution) for further differentiation. 10 µM SU5402 (Sigma-Aldrich; Cat# SML0443-25MG) and 3 µM CHIR99021 (Tocris Bioscience; Cat# 4423) were added to RDM used during the first four media changes. Between Day 75-85 of differentiation, iPSC-RPE cells were purified using magnetic-activated cell sorting as described[88, 89], and the cells were plated on laminin-coated 96-well plates for AAV transduction. Reagent sources were as follows: Heparin (Sigma-Aldrich; Cat# H-3149); ReLeSR (STEMCELL Technologies; Cat# 100-0483); all other differentiation reagents were purchased from ThermoFisher Scientific.

### Viruses and vectors

rAAV2^CBE^ genome was generated by NheI (NEB) enzyme digestion of the Addgene rAAV2 pIC (plasmid infectious clone; cat. 105530-AAV2). Linearized plasmids were isolated from a 1% agarose gel and purified using the GeneJET Gel Extraction Kit (ThermoFischer; K0691). Oligos encoding the AAV2-CBE were designed with single stranded overhangs complementary to NheI (NEB; R3131) generated ends (fwd: CTAGTTGCGACACCATGTGGTCA rev: CTAGTGACCACATGGTGTCGCAA). Oligos were annealed by combining them at equal molar ratios in NEB T4 ligase buffer with T4 Polynucleotide Kinase (NEB; M0201) in a 25µl reaction and incubated at 37°C for 30 minutes followed by boiling at 95°C for 5 minutes and cooled for 1 hour at room temperature. Annealed oligos were ligated and re-circularized into the digested rAAV2 plasmids by T4 DNA ligase (NEB; M0202). DNA was transformed into Stellar^TM^ chemically competent cells (TaKaRa; 636766) and clones were confirmed by sequencing.

wtAAV2 and wtAAV2^ΔCBE^ were made inhouse by co-transfection with Polyethylenimine (PEI). 30ug of DNA at 1:1 molar ratio of pIC:pHelper (expressing Adenovirus E2A, E4, and VA-RNA) were PEI transfected at 3ul/ugDNA into HEK293T cells as described previously[71]. Media was exchanged 5 hours post transfection to avoid toxicity. Cells were incubated for 3 days before cells were collected. Washed cell pellets were lysed by seven freeze-thaw cycles in liquid nitrogen and 37° incubator before being treated with 0.5U/μL DNase I in DNase buffer (Thermo Scientific) for 1hour at 37°C to degrade unencapsidated DNA. Supernatant containing intact capsids was collected through centrifugation at 2,000g for 10min at 4°C. To titer, virus stock was added to TE at a 1:100 dilution and incubated at 57° with Protease Inhibitor Cocktail (MedChem) for 1hour to degrade viral capsids. Viral genomes were quantified by sybr green qPCR with primers complementary to the REP68/78 ORF (fwd: CCGAGAAGGAATGGGAGTT rev: CCATTCCGTCAGAAAGTCG) and compared to corresponding plasmid genomes diluted for a standard curve. rAAV2 and rAAV2^CBE^ vectors for viral titer analysis were produced as above with 30µg of DNA at a 1:1:1 molar ratio of IC:pHelper:pRep2/Cap2. Viral titer was quantified by qPCR with primers complementary to the GFP ORF (fwd: ACGTAAACGGCCACAAGTTC rev: AAGTCGTGCTGCTTCATGTG). For all other experiments rAAV2, rAAV8, and rAAV9 were purchased from Addgene (cat. 105530-AAV2, 105530-AAV8, 105530-AAV9). rAAV2^CBE^, rAAV2/8^CBE^, rAAV2/9^CBE^ virus was produced and provided by Ultragenyx.

### Infection, Transduction, and Transfection

Infections and transductions were performed at the indicated MOI calculated by the number of viral genomes per cell using the viral titers with the doubling time of cells considered. Cells were plated at least 24 hours prior to infection/transduction at which point media was exchanged and the virus/vector was added directly to cells. Plasmid transfections were done at least 6 hours after cells were plated. DNA was transfected by LipoD293™ transfection reagent (SignaGen Laboratories; 50-478-2) at 1µg:3µl ratio. For rAAV2 transduction, >8 weeks old iPSC-RPE cells seeded on 96-well plate were used. The day before transduction, the cells were treated with ViraDuctin^TM^ AAV Transduction reagent (Cell Biolabs; Cat# AAV-201) as per manufacturer’s recommendation. The day of transduction, the cells were washed and treated in triplicates by mixing rAAV2 or rAAV2^CBE^ with 100 µL RDM to get 1E8, E9, or 9E9 vector genome (vg) per well. Untreated cells and cells pre-treated with ViraDuctin^TM^ were used as controls. The next day 125 µL RDM media was added to each well, and on day 2 post-treatment, the media was fully changed. The cells were maintained using regular media changes until flow cytometry analysis.

### CBE in silico analysis

*In silico* analysis of CBEs on viral and vector genomes was performed using JASPAR [90] to screen for all Homo sapiens CTCF consensus sequence profiles using an 80% relative profile score threshold. AAV8 genome sequence was obtained from NCBI (reference sequence NC_006261.1). AAV5 genome sequence was obtained from NCBI (reference sequence NC_006152.1). AAV5 and 8 genomes were aligned to the AAV2 genome, and position of the putative CBE computed relative to the AAV2 reference sequence.

### Western blot

Pelleted cells were lysed on ice for 15 minutes in complete Radio-Immuno-Precipitation Assay (RIPA) buffer (20 mM Tris HCL pH 7.5, 150 mM NaCL, 10% glycerol, 1% NP-40, 1% sodium deoxycholate, 0.1% SDS, 1 mM EDTA, 10 mM trisodium pyrophosphate, 20 mM sodium fluoride, 2 mM sodium orthovanadate and 1x protease inhibitor cocktail (MedChemExpress), glycerol). Cells were centrifuged for 10min at 4°C and 12,000x g and cell lysate was collected. Protein concentrations were calculated using BCA Protein Assay Kit (Bio-Rad) before 6x loading dye was added to final concentration of 1x and samples we boiled at 95°C for 5min. Equivalent protein levels were run on 10% SDS-PAGE gel and transferred to a nitrocellulose membrane. All blocking and antibody incubations were done in 5% milk in TBST (20 mM Tris, 150 mM NaCl, pH 7.6, 0.5% Tween-20) for 45min at room temperature. Membranes were washed in TBST for three 15min washes following 1° and 2° antibodies. Membranes were incubated for 5min with Clarity Western ECL Blotting Substrates (Bio-Rad) and imaged with Li-COR Fortessa scanner.

Antibodies used for western blot assay: CTCF (Cell Signaling; 2899, 1:715), GFP (Cell Signaling; 2955, 1:5,000), and α-tubulin (Millipore; 05-829, 1:5,000). Secondary antibodies HRP-conjugated anti-mouse (Cell Signaling; 7076S, 1:5,000) and anti-rabbit (Cell Signaling; 7074, 1:5,000).

### RT-qPCR

RNA extractions were performed using the Direct-zol RNA Miniprep Kit (Zymo; R2051). Isolated and purified RNA was quantified and 1µg was reverse transcribed using iScript cDNA synthesis kit (Bio-Rad; 1708890) in a 20µl reaction. Resultant cDNA was quantified by sybr green qPCR for 40 cycles. Relative expression was calculated by ΔCT normalized to *Actb* loading control and a no RT reaction control.

Primer complimentary to wtAAV2 REP68/78 ORF (fwd: CCGAGAAGGAATGGGAGTT rev: CCATTCCGTCAGAAAGTCG) or cellular *Actb* (fwd: CACCTTGATCTTCATTGTGCTG rev: GCAAAGACCTGTACGCCAAC) were used.

### RNAi

CTCF knockdowns were performed with two independent siRNAs (S108052: GGAGCCUGCCGUAGAUUtt, S20968: GCUUUGCAGUUACCGUGUtt) or non-specific siMock control (AM4635). Transfections were performed with Lipofectamine RNAiMAX transfection reagent (Thermo Scientific) at a final concentration of 10nM for 24 hours prior to infection with wtAAV2 or wtAAV2^ΔCBE^.

### ChIP-qPCR assays

Cells were crosslinked with 1% formaldehyde for 10 minutes and reaction was quenched in 0.125M glycine for 5min both at room temperature. Cells were scraped, collected, and washed before being lysed in ChIP lysis buffer (1% SDS, 10 mM EDTA, 50 mM Tris-HCl (pH 8), and 1x protease inhibitor cocktail) on ice for 20min. DNA was sheered using Bioruptor for 120 cycles for 30sec on/off. Sonicated supernatant was collected and 10% was frozen for input and remainder was incubated overnight at 4°C with antibody bound protein A Dynabeads™ Protein A (ThermoFischer; 10002D) and ChIP dilution buffer (0.01% SDS, 1.1% Triton X-100, 1.2mM EDTA, 16.7mM Tris-HCl pH 8, 167mM NaCl, 1x protease inhibitor cocktail). Protein bound beads were washed for 3minutes at 4° with low salt wash (0.01% SDS, 1% Triton X-100, 2 mM EDTA, 20 mM Tris-HCl, pH 8, and 150 mM NaCl), high salt wash (0.01% SDS, 1% Triton X-100, 2 mM EDTA, 20 mM Tris-HCl, pH 8, and 500 mM NaCl), lithium chloride wash (0.25 M LiCl, 1% NP-40, 1% deoxycholate, 1 mM EDTA, and 10 mM Tris-HCl, pH 8) and twice with Tris-EDTA (TE) buffer. Samples were eluted in SDS elution buffer (1% SDS, 0.1 M sodium bicarbonate) twice for 15min at 65°C. Cross-linking was reversed using TE buffer and proteinase K (ThermoFischer; EO0491) and incubated for 2 hours at 56°C. ChIP DNA was purified using PCR purification kit (Qiagen) and quantified by Sybr green qPCR analysis (Bio-Rad). Primers complementary to the wtAAV2 REP68/78 ORF (fwd: CCGAGAAGGAATGGGAGTT rev: CCATTCCGTCAGAAAGTCG), rAAV2 GFP ORF (fwd: ACGTAAACGGCCACAAGTTC rev: AAGTCGTGCTGCTTCATGTG).

### Viral Chromosome Conformation Capture coupled with sequencing

Viral Chromosome Conformation Capture (V3C) assay was performed as previously published[68]. Briefly 10 million wtAAV2 or wtAAV2^ΔCBE^ infected HEK293T cells were crosslinked with 1% formaldehyde, and reaction was quenched with glycine. Cells were then lysed and chromatin was digested with HindIII (NEB; R3104) before intramolecular ligation in dilute conditions with T4 DNA ligase (NEB; M0202). DNA was purified before undergoing secondary digest with NlaIII (NEB; R0125) and circularization with T4 DNA ligase under dilute conditions. Viral-host interactions were enriched through inverse and nested inverse PCR amplification steps with primers complementary to wtAAV2 (inverse fwd: CAGATATAAGTGAGCCCAAACG rev: TTCTTGGCTCCACCCTTTT; nested fwd: CATCGACGTCAGACGCGG rev: CTCCACCCTTTTTGACGTAGA). Libraries were prepared using NEBNext Ultra II DNA Library Prep Kit and sequenced using single-end sequencing on an Illumina NextSeq platform spiked with 25% phiX spike-in to increase library complexity. Two independent biological replicates were completed for each sample and processed as below.

Analysis was completed using the Galaxy project server. Sequences were aligned to the hg38 human reference genome using Bowtie2 with default conditions. Aligned sequences were sorted (Samtools sort), formatted (Samtools view) to a BAM format and genome wide coverage was computed using BEDtools[91]. The aligned reads were computed against a library of all HindIII fragments on the hg38 genome using RStudio to generate sum of viral coverage within each HindIII hg38 fragment. The top 150 localization peaks were used in downstream analysis.

Motifs enriched at wtAAV2*ΔCBE* unique sites were determined using the *getfasta* function on BEDtools to extract sequences from the top 150 localization sites, followed by processing using the MEME [92] motif search platform. The transcription factors associated with highly represented motifs were measured using the TOMTOM [92, 93] platform. Intersection analysis was performed using BEDtools Intersect intervals (overlapping either strand, -wa)[91]. Percent localization was computed by SUM((end-start position)*coverage) of intersection results from wtAAV2, wtAAV2^ΔCBE^, or the intersection results of wtAAV2 and wtAAV2^ΔCBE^. Jaccard analysis on BEDtools was used for statistical analysis. The “observed” values were calculated using wtAAV2 and the intersection of wtAAV2 and wtAAV2^ΔCBE^. The “permuted” values were calculated by wtAAV2 intersection with a randomly generated set of genomic coordinates of the same average size and number as wtAAV2-associated regions. Intersections with γH2AX[66], CTCF ChIP-seq[94], and ATAC-seq [95] were specified to direct overlapping interactions. Interactions with histone marks were within a 100bp window. We downloaded the call sets from the ENCODE portal [96] with the following identifiers: ENCSR372WXC, ENCSR372WXC, and ENCSR372WXC.

### Flow Cytometry

HEK293T, U2OS, and HepG2 cells were collected and cell pellets were washed once in PBS prior to being resuspended in PBS with 1ug/ul DAPI live/dead stain (BioRad; 1351303). Cells were immediately put on ice till FACs analysis. Samples were measured on the ThermoFisher Attune NxT V6 Flow Cytometer at the UW-Madison Carbone Cancer Center Flow Cytometry Laboratory. MFI was normalized using standardization Rainbow beads (Spherotech; Cat. RFP-30-5A Rainbow Fluorescent Particles Mid-Range). GFP fluorescence (488nm blue laser) was gated based on the corresponding mock infected cells. Average of 50,000 events were collected per sample. Data was analyzed using FlowJo 10.10.1. Flowcytometry analysis of iPSC-RPE cells was performed as below. To assess transduction efficiency via flow cytometry for GFP expression, cells were washed with 1X DPBS (no Calcium, no magnesium; Thermo Fisher Scientific, Cat # 14190-250) before addition of 1X TrypLE select (ThermoFisher Scientific; Cat # 12563011) for the dissociation of iPSC-RPE monolayer to obtain a single cell suspension. After 30 min incubation at 37°C/5% CO_2_, cells were collected in equal volume of RDM supplemented with 10% FBS and 10 µM ROCK inhibitor Y-27632 (Tocris BioScience; Cat # 1254). Cells were pelleted by centrifugation (400xg, 5 min), resuspended in 400µL of RDM containing 10% FBS and 10 µM ROCK inhibitor, and filtered using Flowmi cell strainers (Bel-Art Products; Cat# H13680-0040) directly in respective pre-labeled round-bottomed tubes (Fisher Scientific; Cat# 14-959-1A) with 1 µg/mL DAPI (Fisher Scientific; Cat# D1306). Flow cytometry was performed using Attune NxT Flow Cytometer (ThermoFisher Scientific) with a 488 nm excitation laser was used to process cells for each sample alongside appropriate controls. For consistency, similar acquisition volume and throughput rates were used for capturing events across all samples. Flow cytometry results were analyzed using FCS Express (De Novo Software).

### CUT&RUN assays

CUT&RUN was performed using the EpiCypher kit (14-1048). 500,000 HEK293T cells infected for 24 hours were used per antibody reaction. Cells were collected and washed with wash buffer before being resuspended in 100µL wash buffer per 500,000 cells and divided into PCR tubes for each reaction. 11ul washed and activated ConA beads were added to cells and incubated 30min at room temperature on nutating mixer. Using a 8-strip magnet to clear bead slurry, supernatant was removed and cell bound beads were resuspended in Antibody buffer with SNAP-CUTANA™ K-MetStat Panel and 1µg antibody of interest. Samples were incubated overnight at 4°C on nutator slightly elevated ensuring bead suspension remains in the bottom of the tube to prevent drying. Using magnet to isolate beads supernatant was removed and beads were washed twice with and then resuspended in 50µL Cell Permeabilization Buffer. pAG-MNase was added to each reaction and incubated for 30 minutes to allow for binding because supernatant was removed and replaced with 200µL Cell Permeabilization Buffer and 1µL 100mM Calcium Chloride to activate digestion for 2 hours at 4°C on nutator before reaction was stopped with a master mix of (0.5ng) E.coli Spike-in DNA and Stop Buffer for 10 minutes at 37°C. Supernatant containing DNA was isolated and purified using SPRIselect beads with 85% ethanol washes and eluted in 15µl TE buffer. Library prep was performed using the EpiCypher kit (14-1001 and 14-1002) Unique pairs of i5 and i7 primers were assigned to each sample and ligated following manufacture protocol with 16 cycles of PCR. DNA was cleaned up using SPRIselect beads and library qualities were verified by Qubit and TapeStation D1000 High Sensitivity tape before pooling samples for sequencing. Samples were pair-end sequenced on a Nextseq2000 (Illumina) at an approximate read-depth of at least 40M read-pairs/sample.

Antibodies used for CUT&RUN from Epicypher: Rabbit IgG (13-0042k), H3K4me3 (13-0060k), H3K27ac (13-0055k), and CTCF (13-2014). Other Antibodies used: H3K9me3 (ab176916).

### CUT&RUN analysis

Analysis of K-MetStat Panel using the Epicypher analysis pipeline determined antibody specificity and overall experimental control (not shown). The resulting paired-end fastq sequencing files were analyzed using the nf-core CUT&RUN pipeline using default settings[97]. Briefly, paired ends were merged and reads underwent adapter and quality trimming (Trim Galore!) before alignment to the specific wtAAV2, wtAAV2^ΔCBE^, rAAV2, or rAAV2^CBE^ genomes (human genome was excluded) using Bowtie2. Reads were sorted, and quality checked by samtools and picard before processed into coverage files (BEDtools) which were used for downstream processing. Genome coverage was computed from the aligned BAM files and scaled to sequencing depth using reads per million mapped reads (RPM). This normalization was performed independently for each dataset to ensure comparability across samples with different read depths. Samples were then normalized to corresponding IgG control signal per bin across the genome. Normalized RPM signal was calculated by (Target RPM) – (IgG RPM). Negative values were set to zero and signal peaks over IgG levels were further processed. IgG normalization was performed by matching each target replicate to its corresponding IgG control replicate. However, for a single replicate of wtAAV2^ΔCBE^ H3K4me3, H3K27ac, and CTCF, the experimental IgG control failed to produce a bedGraph file due to a processing error. To evaluate its suitability for inclusion, the IgG BAM file from this replicate was compared to the corresponding IgG BAM files from the other replicates, yielding strong correlations (Pearson r ≥ 0.8). Given this concordance, normalization was performed by pairing each target replicate with all available IgG replicates. Target signals were normalized independently to each IgG control, and the resulting normalized values were averaged on a per-bin basis. The averaged normalized signal was used for downstream analyses. Replicate normalized RPM peaks were averaged at each base pair and plotted on UCSC tracks on the respective genomes. Quantification of mean RPM/bp was calculated by SUM[((end-start position)*normalized count)/genome size] of individual replicates. ITR region was removed from whole genome quantification.

### Statistical analysis

Statistical analysis was performed using GraphPad Prism software and appropriate statistical tests were done with t-tests, Ordinary one-way Anova, or Ordinary two-way ANOVA followed by either Dunnett’s multiple comparisons test or Tukey’s multiple comparisons test.

## DATA ACCESSION

All high-throughput sequencing data has been uploaded to the NCBI’s public archive at Gene Expression Omnibus (GEO). The CUT&RUN data is available under the accession number GSE334008, wtAAV2^CTCF^ V3C-seq under the accession number GSE334009 and the wtAAV2 V3C-seq under the accession number GSE320210.

## ACKNOWLEDGEMENTS

This research was funded partially by NIH/NIAID K99/R00 Pathway to Independence Award, grant number AI148511, to K.M.; the Wisconsin Partnership Program’s New Investigator Award (PERC Grant G-4942) to K.M. and NIH/NIGMS R35 Maximizing Investigator’s Research Award (MIRA), grant number GM154938, to K.M. C.I.S.L. is funded by NSF Graduate Research Fellowship Program award DGE-2137424. R.R.A. was funded by a SciMED Graduate Research Scholarship from the University of Wisconsin-Madison and NIH F31 Ruth L. Kirschstein Pre-Doctoral Fellowship (AI191692). We thank the Precision Medicine Research Service of the UW Center for Human Genomics and Precision Medicine for high-throughput sequencing. This work was supported in part by a Sponsored Research Agreement from Ultragenyx Pharmaceuticals, to K.M.; an Unrestricted Grant from Research to Prevent Blindness, Inc. to the UW-Madison Department of Ophthalmology and Visual Sciences, the Sandra Lemke Trout Chair in Eye Research (DMG), the Retina Research Foundation Emmett A. Humble Distinguished Directorship of the McPherson Eye Research Institute (DMG), a core grant to the Waisman Center from the National Institute of Child Health and Human Development (P50HD105353), an unrestricted grant from Research to Prevent Blindness to the UW-Madison Department of Ophthalmology and Visual Sciences, and University of Wisconsin Carbone Cancer Center Flow Cytometry Laboratory (supported by P30CA014520). We acknowledge Dr. Sara Maloney (IMV, UW Madison) and Dr. Kavi Mehta (CBMS, UW Madison) for critical review of the manuscript and scientific discussions.

---

**Supplemental Figure S1:**
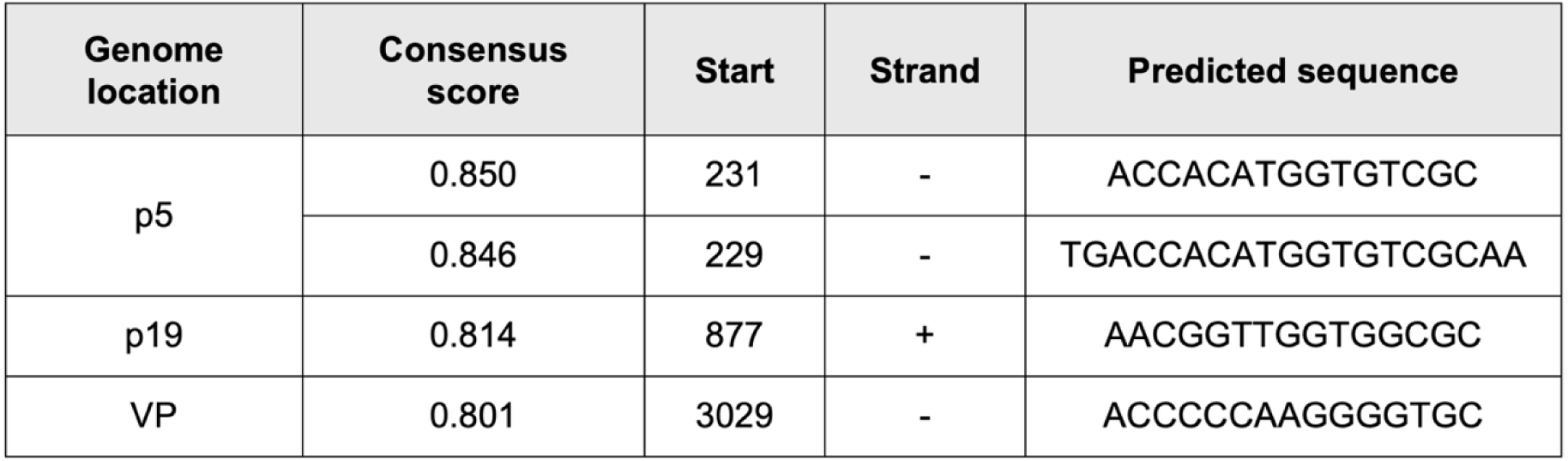
wtAAV2 *in silico* CBEs. Results of JASPAR informatics resource *in silico* prediction analysis of CBEs on wtAAV2. Table shows the position and strand. Consensus score is calculated by Position-Specific Weight Matrix (PWM) (score ≥ 0.8 threshold for a strong candidate binding site).

**Supplemental Figure S2:**
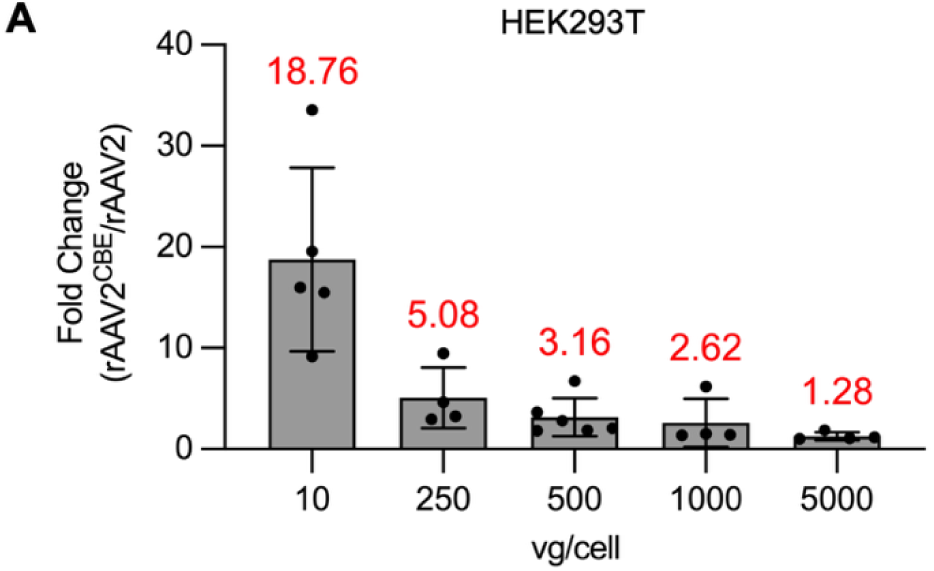
Comparison of rAAV2 and rAAV2^CBE^ transduction efficiencies. A) Fold change of percentage of GFP positive single cells from flow cytometry analysis of rAAV2^CBE^ relative to rAAV2 at indicated MOI. Fold change shown in red. Dots represent individual calculated values with bar showing mean and error bars represent standard error mean (SEM).

**Supplemental Figure S3:**
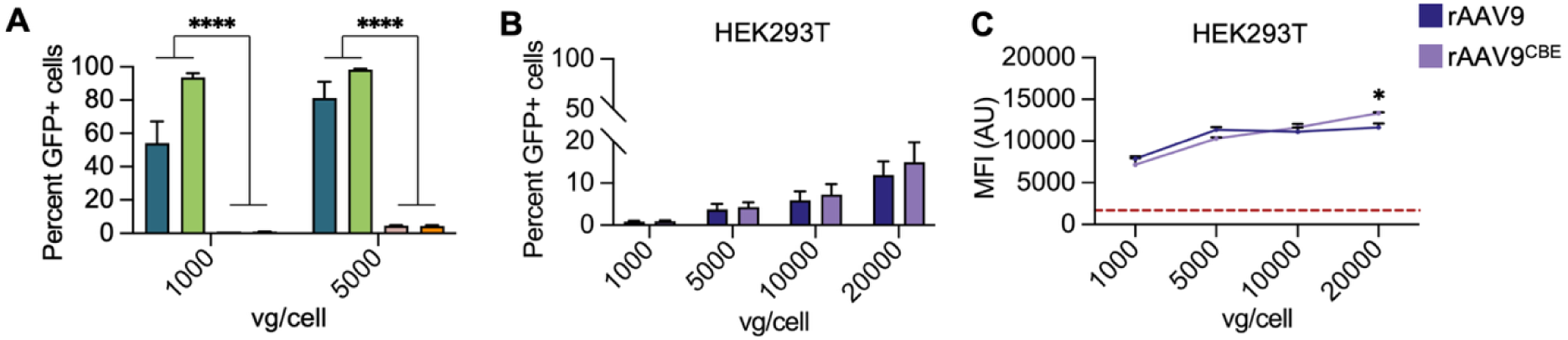
AAV2-CBE in rAAV8 and rAAV9. A) Comparison of rAAV2 and rAAV2/8 transductions in HEK293T cells at indicated MOI. rAAV2 (teal), rAAV2^CTCF^ (green), rAAV2/8 (pink), rAAV2/8^CTCF^ (orange). B) Quantification of flowcytometry percentage of single cells that are GFP positive over increasing MOI comparing rAAV2/9 (dark purple) and rAAV2/9^CTCF^ (light purple). G) Median Fluorescent Intensity of GFP in arbitrary units from flowcytometry over increasing MOI. Red dashed line represents average mock MFI. HEK293T cells 24 hpt (n=3). Error bars represent standard error mean (SEM). Statistic *p*-value ns (not shown) >0.5, * <0.03, ** <0.002, *** <0.0002, **** <0.0001.

